# Using set theory to reduce redundancy in pathway sets

**DOI:** 10.1101/319731

**Authors:** Ruth Stoney, Jean-Mark Schwartz, David L Robertson, Goran Nenadic

## Abstract

1.

**Background:** The consolidation of pathway databases, such as KEGG[1], Reactome[2]and ConsensusPathDB[3], has generated widespread biological interest, however the issue of pathway redundancy impedes the use of these consolidated datasets. Attempts to reduce this redundancy have focused on visualizing pathway overlap or merging pathways, but the resulting pathways may be of heterogeneous sizes and cover multiple biological functions. Efforts have also been made to deal with redundancy in pathway data by consolidating enriched pathways into a number of clusters or concepts. We present an alternative approach, which generates pathway subsets capable of covering all of genes presented within either pathway databases or enrichment results, generating substantial reductions in redundancy.

**Results:** We propose a method that uses set cover to reduce pathway redundancy, without merging pathways. The proposed approach considers three objectives: removal of pathway redundancy, controlling pathway size and coverage of the gene set. By applying set cover to the ConsensusPathDB dataset we were able to produce a reduced set of pathways, representing 100% of the genes in the original data set with 74% less redundancy, or 95% of the genes with 88% less redundancy. We also developed an algorithm to simplify enrichment data and applied it to a set of enriched osteoarthritis pathways, revealing that within the top ten pathways, five were redundant subsets of more enriched pathways. Applying set cover to the enrichment results removed these redundant pathways allowing more informative pathways to take their place.

**Conclusion:** Our method provides an alternative approach for handling pathway redundancy, while ensuring that the pathways are of homogeneous size and gene coverage is maximised. Pathways are not altered from their original form, allowing biological knowledge regarding the data set to be directly applicable. We demonstrate the ability of the algorithms to prioritise redundancy reduction, pathway size control or gene set coverage. The application of set cover to pathway enrichment results produces an optimised summary of the pathways that best represent the differentially regulated gene set.

## 2. Background

Pathways are sets of genes corresponding to functionally related interacting proteins. Pathway data is available from many databases dependent on biological focus. The fragmented nature of pathways across multiple databases makes it difficult to perform inclusive analysis of all known data. To address this issue, many attempts have been made to consolidate pathway databases such as ConsensusPathDB (CPDB) [4], PathwayCommons [5], The Human Pathway Database (HPD) [6], Pathway Interaction Database (PID) [7], and NCBI Biosystems [8]. Amalgamating multiple databases into a consistent searchable format facilitates the use of these resources, however the arbitrary nature of pathway boundaries results in overlap and redundancy. This redundancy greatly increases the quantity and complexity of pathway data, which has lead to the development of a range of tools to assist in data simplification and interpretation [6, 7, 9–11]. Previous solutions presented to deal with redundancy include visualizing redundancy between pathways to the user [6], merging pathways based on similarity [10, 11] and even integrating full pathway sets into a non-redundant, single unified pathway [12]. Reducing redundancy simplifies the pathway-related descriptive space, allowing multiple resources to be combined while limiting the number of pathway attributes assigned to each gene. The advantages are apparent, with resources such as PathCards being integrated into the widely used GeneCards[11].

Redundancy Control in Pathway Databases (ReCiPa) [10] uses a pathway merging algorithm to combine pathways with high levels of overlap. Users select a maximum overlap threshold and pathway pairs displaying greater levels of overlap are merged. Within that study redundancy was observed within five large databases (KEGG, Biocarta, CGP, NCI-PID, and Reactome). They proceeded to merge pathways from the Molecular Signatures Database (MSigDB), whose overlap exceeded 75%, reducing pathway redundancy.

Pathcards described a multistep procedure to reduce pathway redundancy, also through pathway merging [11]. Two thresholds were calculated and sequential merging steps were used to minimize overlap, while preventing the generated super-pathways from becoming too large to be informative. By merging pathways into super-pathways, Pathcards suggested many new molecular interactions. They demonstrated that many of these newly generated interactions are supported by high numbers of literature co-mentions and high experimental interactions scores according to STRING. However, while the generation of potential interactions can be highly beneficial, if the aim is to utilize previously validated data, merging pathways introduces a source of uncertainly into the dataset.

A major application of pathway data sets is pathway enrichment analysis. Both Pathcards and ReCiPa explored the capability of their reduced pathway dataset to improve enrichment results. Enrichment analysis of 830 differential expression sets was performed using the super-pathways generated within Pathcards. The enrichment results from super-pathways tended to be more significant than the enrichment scores of their constituent pathways. Similarly within the ReCiPa study enrichment analysis was performed using genes differentially expressed in obesity. After merging, the top 20 most significantly enriched pathways showed less overlap and greater significance towards the disease, compared to the original dataset.

Pathway Distiller implemented an alternative approach by removing redundancy from enriched pathway sets following enrichment analysis [9]. Pathways may be consolidated into pathway concepts based on gene expression profiles, gene membership, protein-protein interaction data or shared Gene Ontology (GO) terms. Each method provides varying, complementary views of the data, with different pathway concepts generated. Consolidating enrichment output into a reduced number of pathway concepts increases data manageability and readability, by organizing redundant pathways into their major groups.

All of the approaches discussed to this point have used merging and consolidation to address redundancy. Alexa et al. (2006) demonstrated that redundancy in GO enrichment results could be reduced by selecting a subset of representative terms [13]. Pathway enrichment analysis and GO enrichment analysis are similar techniques in which sets of differentially expressed genes are compared to gene sets associated with pathways or GO terms. Alexa et al. (2006) introduced two algorithms, *elim* and *weight*, which use the Gene Ontology topology to select a representative subset of highly enriched GO terms [13]. The enrichment set cover algorithm presented in this paper shares some conceptual similarity with this approach however, the implementation is different since there is no organized topological hierarchy for combined pathway datasets and the rules governing the Gene Ontology, such as the true path rule [14], do not apply.

Within this paper we show that set cover can be used to reducing redundancy by selecting subsets of representative pathways. We describe a set of algorithms for reducing redundancy in pathway datasets, as well as a separate algorithm for reducing redundancy from pathway enrichment results. The proportional set cover algorithm and hitting set cover algorithm aim to identify a minimum subset of pathways required to cover the genes in highly redundant, consolidated pathway databases. The generated set covers are not designed to depict the full range of possible pathway boundaries and their accompanying cellular functions, but rather they provide a simplified set of pathways to represent the actions of genes within the dataset. Since the pathways are not merged database and biological information remains directly applicable and functional specificity is not lost through pathway size expansion. The proposed method also removes the risk of biologically distinct pathways being merged. The algorithm’s ability to remove overlap is not limited by thresholds, conferring an advantage compared to approaches such as Pathcards and ReCiPa in which redundancy between pathway pairs can only be removed if the overlap exceeds the threshold. Set cover algorithms also consider redundancy between multiple pathways, rather than just comparing pathway pairs.

We also developed the enrichment set cover algorithm for handling pathway enrichment data and applied it to a set of enriched osteoarthritis pathways [15]. In contrast to the approaches used by ReCiPa and Pathcards, the enrichment set cover algorithm is designed to be used following enrichment analysis, which should be performed using the full pathway dataset. Redundancy is then removed from the enriched pathway set by selecting the pathway with the lowest p-value to cover each differentially regulated gene. Enriched pathways are not merged or altered and the number of enriched pathways required to cover the dataset is reduced. The resulting pathways set can therefore be used as an optimized summary output, conveniently showing the most important pathways for describing the differentially regulated gene set. By increasing the number of differentially regulated genes covered by the most highly enriched pathways, researchers examining the top 10 or 20 pathways are provided with a more inclusive portrayal of the gene set.

## 3. Approach

We downloaded pathway data from ConsensusPathDB (CPDB), an opensource online collection of pathways, that incorporates 32 sources including KEGG, Wikipathways, PDB, Reactome. CPDB makes these resources available as a single download, which we acquired on 24/09/2015 containing 4,011 pathways. We applied the set cover algorithm to the CPDB data set, analyzing it’s effectiveness at: reducing pathway overlap; reducing pathway size variability; and preserving the maximum number of genes in the data set. We found that standard set cover caused unacceptable increases in pathway size, therefore we modified the algorithm and assessed the modified algorithms capability to meet the previous three objectives.

Set cover is a well-defined algorithm in computer science for handling overlapping sets of sets. For example, set cover is used by CLASS, a bioinformatics program that maps RNA sequence data to transcripts [16]. Set cover has also been used to predict protein-protein interactions based on binding domains [17], to reduce the complexity of SNP sets [18] and to minimize the number of probes needed to analyze DNA [19].

Set cover algorithms deal with elements and sets, which relate to genes and pathways respectively. All the unique genes in the data set are collectively referred to as the universe. The aim is to produce a reduced selection of sets (pathways), which collectively cover all the elements (genes) in the universe (dataset). This subset of the original data is called the cover set [20]. Each time a pathway is added to the cover set the genes in the pathway become covered (Figure 1). Direct application of set cover lead to extremely large, functionally non-specific pathways dominating the cover set, therefore we implemented the proportional set cover and hitting set cover algorithms to better control pathway size, while reducing redundancy and covering the dataset.

**Figure 1.**
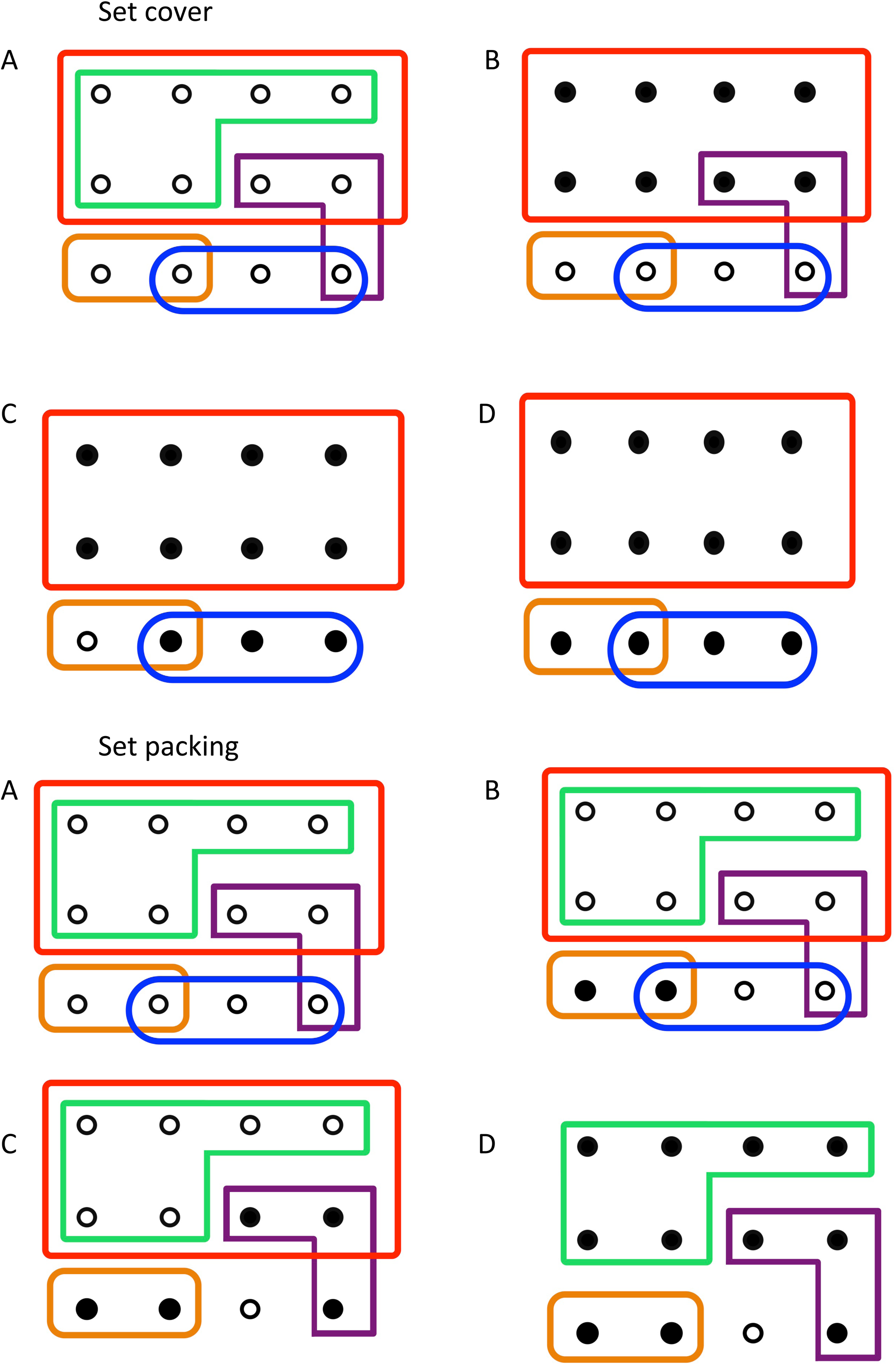
Set cover A) A simple set of overlapping sets. B) The red set with 8 uncovered elements is selected first. C) The blue set with 3 elements is selected second. D) The orange set then covers all the elements in the universe.

When dealing with enrichment analysis data the aim is to reduce redundancy between pathways, while preserving the order of enrichment significance denoted by the p-values. We designed an algorithm that would select the set of pathways with the lowest p-values capable of covering all the genes in the dataset. This ensures that the filtered results return the most enriched pathways available for each gene.

## 4. Methods

### 4.01 Overlap score

To measure overlap across different algorithms we measured the mean number of pathways that each gene appears in. Within the raw data genes appeared in a mean of 12.4 pathways. We refer to this metric as the overlap score.

### 4.02 Set cover

We applied the set cover algorithm to the data set, which generates a subset of pathways called a cover set, in which all the genes in the data set are represented or “covered”. Set cover begins by first assigning values to each pathway (*v_i_*). The set cover values correspond to the number of uncovered genes each pathway contains (Equation 1).

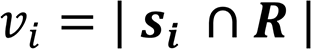

where (***s****_i_*) is the pathway’s gene set and ***R*** is the set of all uncovered genes.

At the beginning of the algorithm all the genes in the dataset are uncovered so the algorithm selects the largest pathway. The genes from the selected pathway are then covered, so it is unnecessary to cover them again using additional pathways. The algorithm then recalculates how many uncovered genes each pathway contains and continues to add the pathway with the maximum value to the set cover until all genes in the data set are covered.

#### Algorithm 1

Set cover (in separate file)

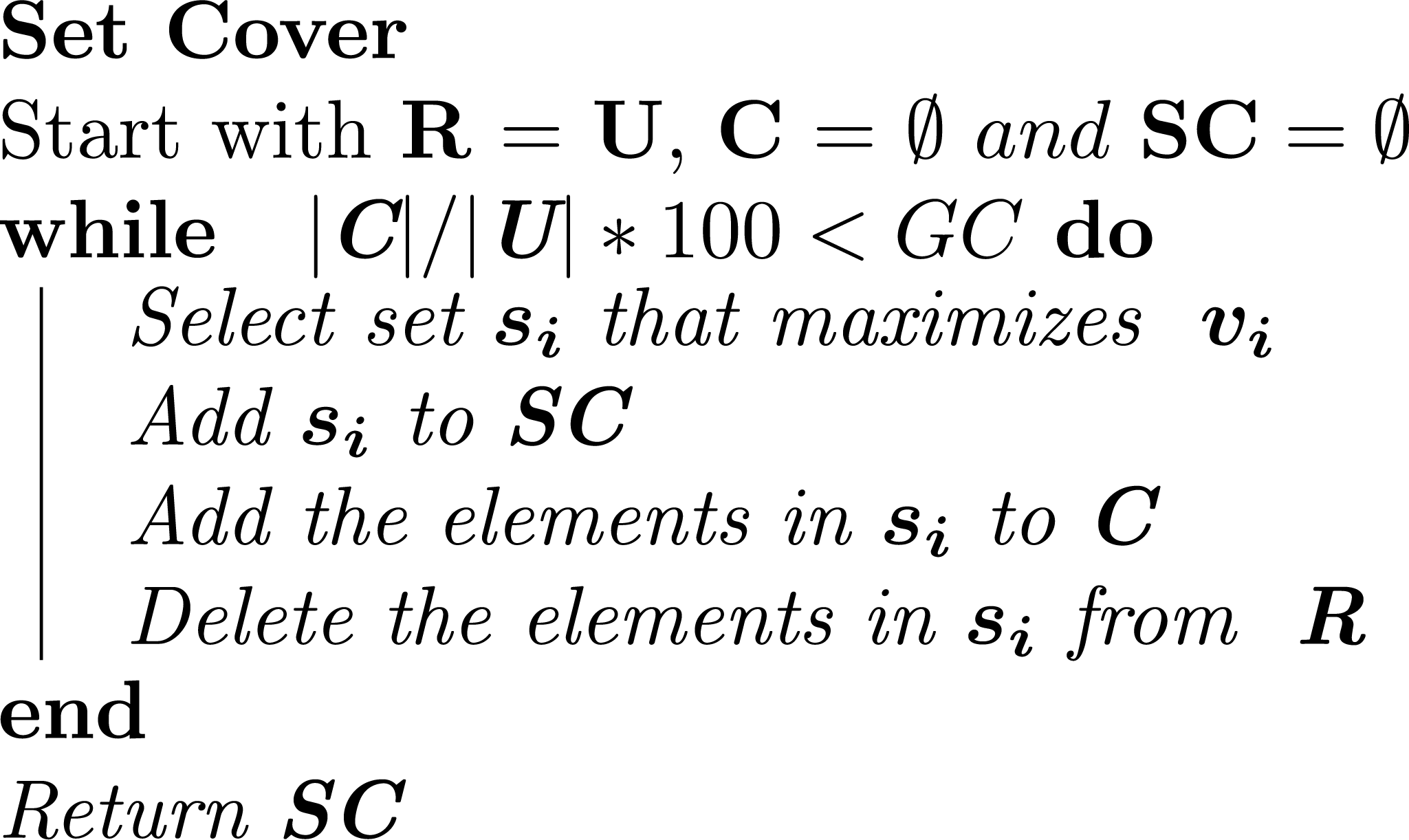

where ***R*** is the set of uncovered genes, ***U*** is all the genes in the dataset, ***C*** is the covered genes, ***SC*** is the set cover result, *GC* is the gene coverage (see Section 4.03) and ***s_i_*** is a pathway.

Application of the set cover algorithm was effective in reducing overlap between the pathways; however, it selected very large pathways with reduced informativeness (maximum size 2320, standard deviation 160, almost double the standard deviation on the original dataset 86.9). We therefore explored methods that avoid preferential selection of large pathways.

### 4.03 Gene Set Coverage

As the set cover algorithm approaches completion and the final sets are added to the cover set, increases in data coverage are gained at the expense of redundancy reduction. This is because the final sets required to cover the few remaining genes tend to have the most overlap with other pathways already in the set cover. In addition, fewer pathways are available to cover the final few genes, restricting options to control pathway size. To allow a user-defined compromise between the gene coverage, pathway redundancy and pathway size we introduce the Gene Coverage (*GC*) parameter. Setting *GC* below 100% allows the algorithm to finish before the final elements have been covered. We experimented setting *GC* to 90, 95, 99 and 100% of the number of genes in the data set.

### 4.04 Proportional set cover

When reducing pathway redundancy there are three competing aims: reducing redundancy; controlling pathway size; and covering the entire gene set. The proportional set cover algorithm was generated to focus on controlling pathway size.

To control the size of the pathways we altered the scoring mechanism to rank pathways based on the proportion of uncovered genes they contained, rather than the absolute number (Equation 2). This works because larger pathways are more likely to have a proportion of their genes covered when other pathways are selected. Additionally this mechanism directly penalizes overlap, which the standard algorithm does not. At the beginning of the proportional set cover algorithm none of the genes are covered so the proportion of uncovered genes in every pathway is 1. This would result in the starting pathway being selected at random. To ensure that pathway size variability is controlled as strictly as possible, we implemented the second part of Equation 2, which ensures that pathways of mean pathway size are preferentially selected when multiple pathways with the same proportion of uncovered genes are available.

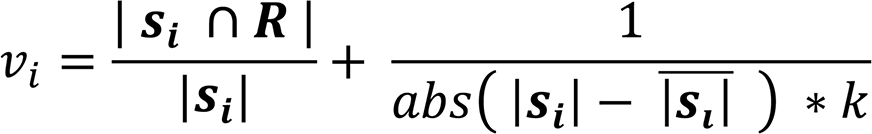

where ***s****_i_* is the pathway’s gene set, 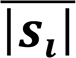 is the mean pathway length, ***R*** is the uncovered genes set and *k* is a large constant to limit the influence of the second term (taken equal to 10,000).

### 4.05 Hitting set cover

The set-covering problem can be reformulated into the equivalent set-hitting problem. In this formulation genes and pathways are visualized as bi-partite graph in which the pathways are connected to the genes that they contain. In this depiction it is clear that some genes are only linked to a single pathway, which must be selected if the gene is to be covered. The importance of pathways can therefore be considered as a factor of how infrequent their genes are. The hitting set cover is therefore designed to reduce redundancy as much as possible without directly selecting for pathway size.

We calculated the frequency of each gene in the data set (*F*), then assigned the gene’s value *gv(j)* as 1/*F*. We then assigned a value *v_i_* to each pathway defined as the sum of each uncovered gene’s scores divided by the number of genes in the pathway (Equation 3).

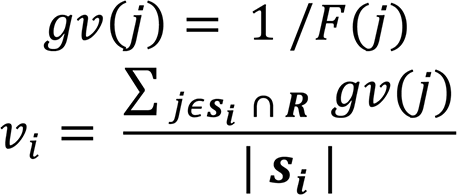

where *gv(j)* is the value of a gene, *F(j)* is the number of pathways a gene is in, *jϵ****s****_i_* ∩ ***R*** means for each uncovered gene in the pathway and |***s****_i_*| is the length of the pathway.

### 4.06 Set cover for pathway enrichment analysis

Pathway analysis is a frequently used method; therefore a modified set cover algorithm to address this situation could be highly useful. The universe represents differentially expressed genes and the sets are enriched pathways generated through enrichment analysis. Enrichment analysis results represent entirely different input data compared to the pathway datasets used in the previous algorithms, as the enriched pathways already have scores (p-values). We wish to reduce redundancy (gene overlap) between enriched pathways and it is essential that the pathways with the lowest possible p-values are selected. Equation 4 allows the pathways with the lowest p-values to be selected, unless all of their genes are covered by other enriched pathways with even lower p-values.

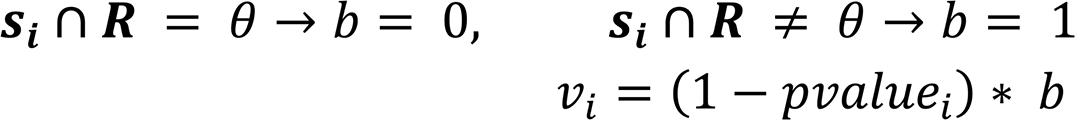

where ***s****_i_* is the enriched pathway’s gene set, ***R*** is the uncovered gene set, *b* is a binomial operator, *pvalue_i_* is the pathway’s p-value and *v_i_* is the pathway’s set cover value.

We generated the enriched data set by applying GOseq [21] to expression data from the damaged cartilage in osteoarthritis patients and controls [15].

## 5. Results

We started with the large, extensively redundant CPDB data set and used set cover to reduce pathway overlap, while controlling pathway size and seeking to cover as much of the data set as possible. We describe the ability of the standard set cover algorithm and two modified algorithms, in conjunction with the *GC* parameter, to meet these objectives.

### 5.01 Pathway redundancy varies between different algorithms

The original pathway data set contained 11,196 genes and 3,305 pathways; the starting overlap score (see methods) was 12.4. The standard set cover algorithm reduced overall redundancy from 12.4 to 4.1, a 73% reduction (since a completely discrete pathway set would have a score of 1). The overlap score for proportional set cover was 4.36, slightly higher than the standard set cover algorithm, but still representing a 70% reduction in overlap from the original data. The hitting set cover algorithm was designed to select pathways that contained rare genes within the data set, resulting in the greatest reduction in overlap (overlap score of 3.95 equivalent to a 74% reduction).

After application of the set cover algorithms the distribution of the remaining overlap between pathways varied greatly. Figure 2 shows the Jaccard similarity between pairs of pathways, in the outputs produced by each of the three algorithms. The standard set cover algorithm produced the lowest maximum overlap (Jaccard similarity = 0.68) between the pathway pairs. However, compared to the original data, a higher proportion of pathway pairs in the set cover output showed Jaccard similarities between 10–30%. Proportional set cover had the greatest maximum Jaccard similarity at 0.93, out of the set cover algorithms. The hitting set cover algorithm produced a maximum Jaccard similarity between two pathways of 0.82, despite having the lowest overlap score.

**Figure 2.**
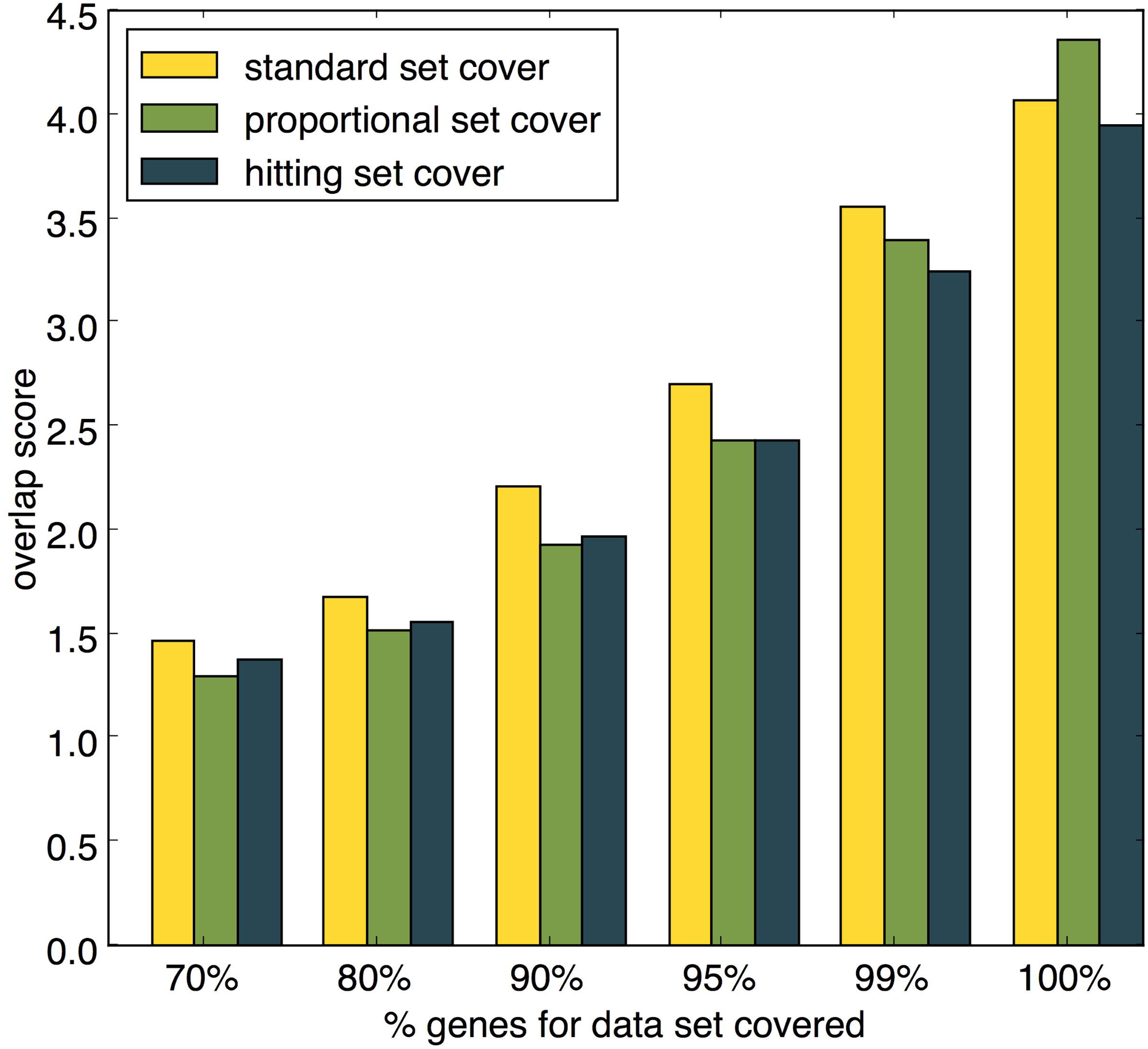
Jaccard coefficient between pathway pairs in the cover set results produced by each algorithm.

#### Gene Coverage can be lowered to reduce redundancy

For each of the algorithms it is possible to use the *GC* parameter to prioritize reductions in redundancy over gene coverage by stopping any algorithm before all of the genes in the dataset have been covered. Figure 3 shows improved ability of the set cover algorithms to reduce pathway overlap for different values of *GC*. If 99% of the genes are required then the hitting set algorithm achieves the lowest overlap score of 3.24, equivalent to an 80% reduction in overlap. Redundancy can be further reduced if only 95% of the genes are covered, with the proportional and hitting set algorithms producing an overlap score of 2.41, equivalent to a 88% reduction in redundancy. Both the proportional set cover and the hitting set cover are more effective at reducing redundancy than the standard set cover if *GC* is set to less than 100%.

**Figure 3.**
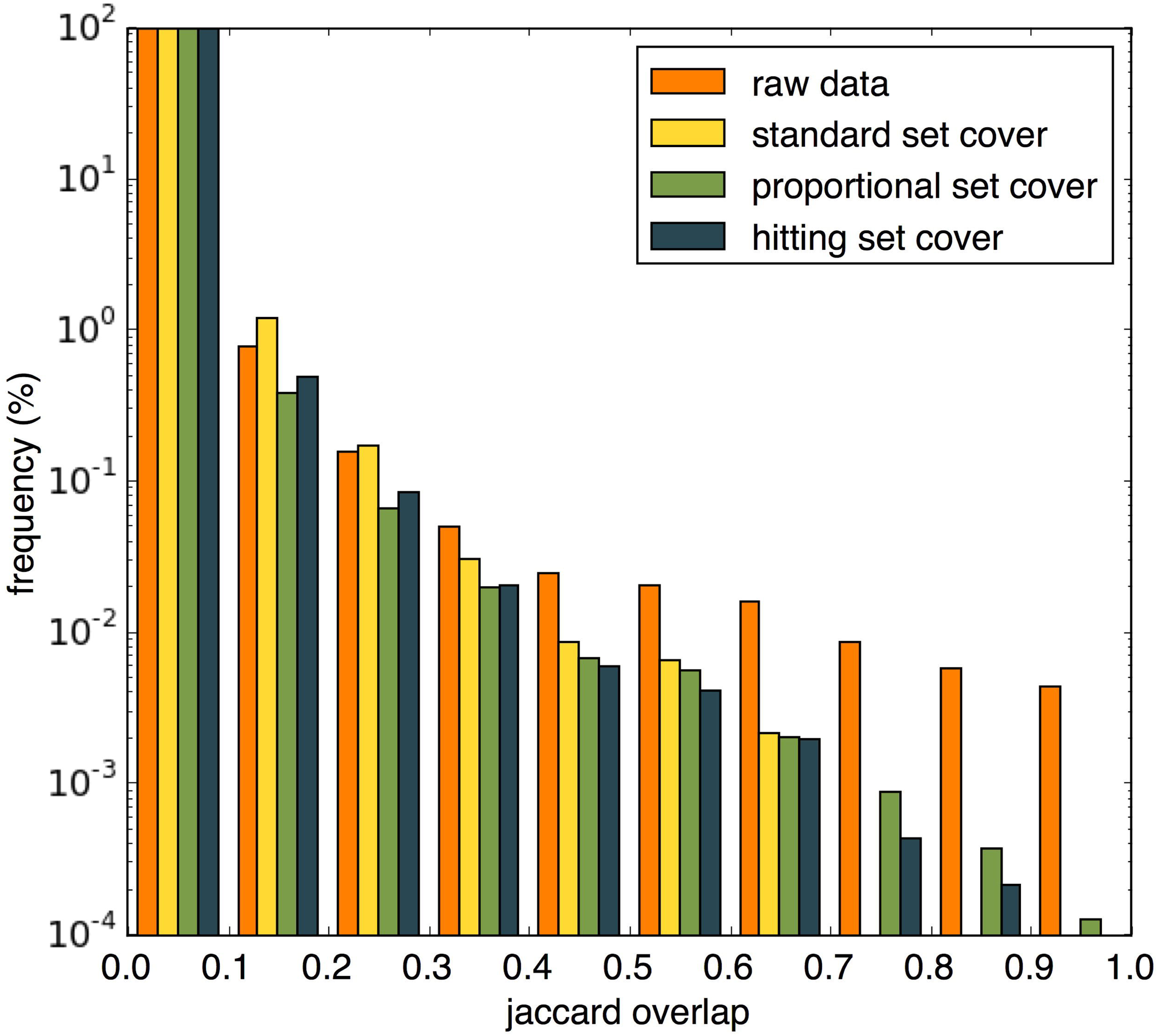
Redundancy in set cover outputs given different GC values.

#### Pathway size is affected by the set cover algorithm and Gene Coverage setting

When *GC* was set to 100% the standard set cover algorithm represented all of the genes in the dataset using only 524 pathways (16% of the original pathway set). However, many of these were very large increasing the mean size to 87.2 (standard deviation 160.1). These pathways have reduced informativeness since functional specificity is lost. Figure 4A illustrates the tendency of this algorithm to select extremely large pathways.

**Figure 4.**
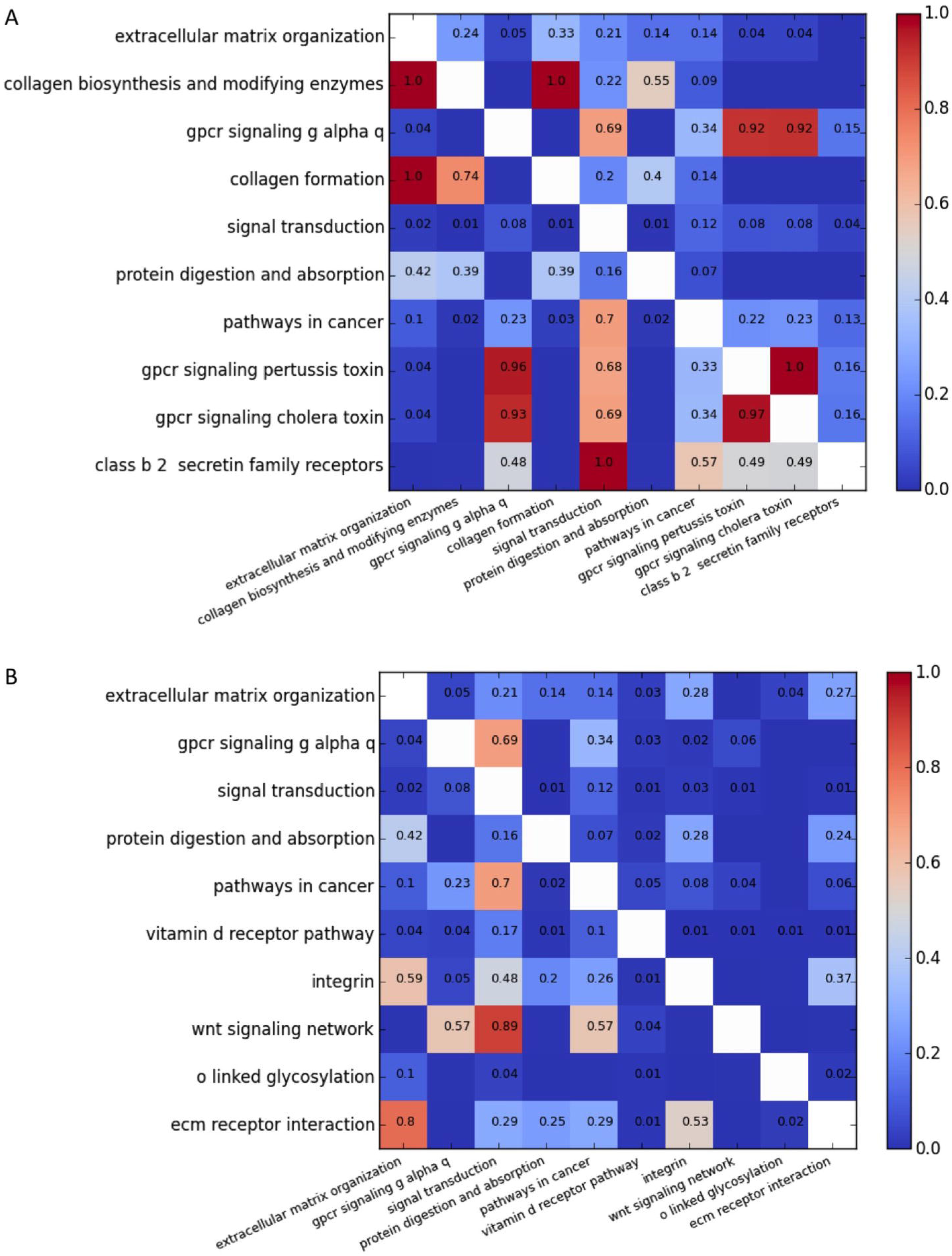
Pathway sizes in cover set when GC is set to A) 100%, B) 99%, C) 95% and D) 90%. The boxes indicate the 25th and 75th percentiles and the whiskers indicate the 5th and 95th percentiles.

The proportional set cover algorithm was designed to preferentially select moderately sized pathways. This returned a cover set of 1,336 pathways with controlled size variation (mean of 36.5, standard deviation 55.1) shown in Figure 4A. The hitting set cover algorithm was less able to control pathway size than the proportional set cover algorithm, returning 957 pathways with a mean size of 46.2 (standard deviation 61.7).

Figures 4B – D show that as *GC* is reduced the tendency of the standard set cover to select very large pathways becomes more exaggerated. Decreasing *GC* also improves the ability of the proportional set cover algorithm to select moderately sized pathways. The hitting set algorithm also tends to select smaller pathways when *GC* is reduced, since larger pathways often contain more frequent genes. Reducing *GC* affects pathway size since in the later stages of the algorithm, fewer pathways are available to cover the remaining genes, reducing the available options. Therefore, lowering *GC* has the ability to help control pathway size when the proportional set cover and hitting set cover algorithms are used.

*Since the databases that contribute to CPDB contain pathways of different sizes, the set cover generated may preferentially select pathways from some databases more than others*.

Table 1 shows the proportion of pathways that come from each database in the cover set generated by each algorithm. All algorithms generate set covers with reduced INOH and SMPDB pathways, showing that SMPBD’s focus on small molecules and INOH’s ontology-based approach tend to be ill-suited to the generation of discrete pathway protein sets. The standard set cover algorithm generates sets containing large pathways, preferentially selecting pathways from KEGG (median size 65, see Table 1) and Netpath (median size 51); while proportional set cover tends to select smaller pathways from Reactome (median size 17), HumanCyc (median size 5) and Signalink (median size 32), whilst avoiding NetPath.

**Table 1.** Proportion of pathways from CPDB databases. Median size represents the median sizes of the pathways in the CPDB dataset. CPDB % represents the proportion of the pathways in the unaltered dataset that came from each database. The following columns represent the proportion of pathways in the set cover generated by the standard set cover algorithm, the hitting set cover algorithm and the proportional set cover algorithm. Different results are obtained by altering the proportion of the gene set covered, shown in subcolumns below the algorithm header.

|  | Median size | CPDB % | Standard set cover |  |  |  | Hitting set cover |  |  |
| --- | --- | --- | --- | --- | --- | --- | --- | --- | --- |
|  |  |  | 100% | 99% | 95% | 90% | 100% | 99% | 95% |
| BioCarta | 15.0 | 6.3 | 6.3 | 4.6 | 0.5 | 0.0 | 4.7 | 4.8 | 5.4 |
| EHMN | 32.5 | 1.6 | 3.2 | 3.4 | 2.6 | 1.0 | 2.1 | 2.3 | 1.8 |
| HumanCyc | 5.0 | 8.2 | 6.5 | 7.7 | 2.6 | 0.0 | 10.1 | 10.9 | 12.9 |
| INOH | 34.5 | 2.3 | 1.7 | 1.9 | 1.0 | 1.0 | 0.8 | 0.6 | 0.3 |
| KEGG | 65.0 | 7.2 | 29.0 | 30.5 | 37.6 | 40.4 | 15.8 | 15.0 | 13.5 |
| NetPath | 51.0 | 0.9 | 2.1 | 2.4 | 3.6 | 5.1 | 1.1 | 1.2 | 1.1 |
| PharmGKB | 13.0 | 2.8 | 3.1 | 2.9 | 0.5 | 0.0 | 2.0 | 2.1 | 2.4 |
| PID | 35.0 | 5.2 | 15.6 | 13.9 | 10.3 | 6.1 | 9.5 | 9.8 | 9.4 |
| Reactome | 17.0 | 39.6 | 4.2 | 5.3 | 10.8 | 21.2 | 36.1 | 35.1 | 34.7 |
| Signalink | 32.0 | 0.4 | 1.0 | 1.2 | 1.0 | 0.0 | 0.6 | 0.7 | 0.7 |
| SMPDB | 11.0 | 16.7 | 1.7 | 1.4 | 0.5 | 0.0 | 1.6 | 1.5 | 1.4 |
| Wikipathways | 26.0 | 8.8 | 25.6 | 24.9 | 28.9 | 25.3 | 15.6 | 16.0 | 16.2 |

#### Reducing redundancy in pathway enrichment analysis

To demonstrate the ability of the set cover algorithm to handle enrichment data, we applied the enrichment set cover algorithm to an osteoarthritis data set, retrieved from Dunn et al. (2016) [15]. From the osteoarthritis data set, 58.3% of the differentially expressed genes could be mapped to a CPDB pathway, which was a 17% improvement on the GOseq [21] implemented data set. We retrieved 42 enriched pathways with a p-value lower than 0.05, following the Benjamini-Hochberg correction for multiple testing. Set cover for enrichment analysis reduced the number of pathways required to cover the differentially expressed genes to 23 (supplementary table 1).

The heat map in Figure 5A shows the asymmetric overlap between the top ten pathways before application of the algorithm. The p-values from pathway enrichment determine the order in which pathways were considered for inclusion in the cover set. Pathways were omitted if all of the differentially expressed genes that they covered were also covered by more enriched pathways. Note that overlap tends to be higher in the bottom left triangle as pathways added later were often smaller subcomponents of larger pathways. We can see that ‘extracellular matrix organization’, the most enriched pathway, was placed in the cover set first. Next was ‘collagen biosynthesis and modifying enzymes’; however, all of the differentially expressed genes in this pathway are also covered by the larger pathway ‘extracellular matrix organization’, as indicated by the red cell in the ‘collagen biosynthesis and modifying enzymes’ row, ‘extracellular matrix organization’ column. The corresponding cell in the ‘extracellular matrix organization’ row reveals that 24% of the differentially expressed genes in ‘extracellular matrix organization’ are also in ‘collagen biosynthesis and modifying enzymes’.

**Figure 5.**
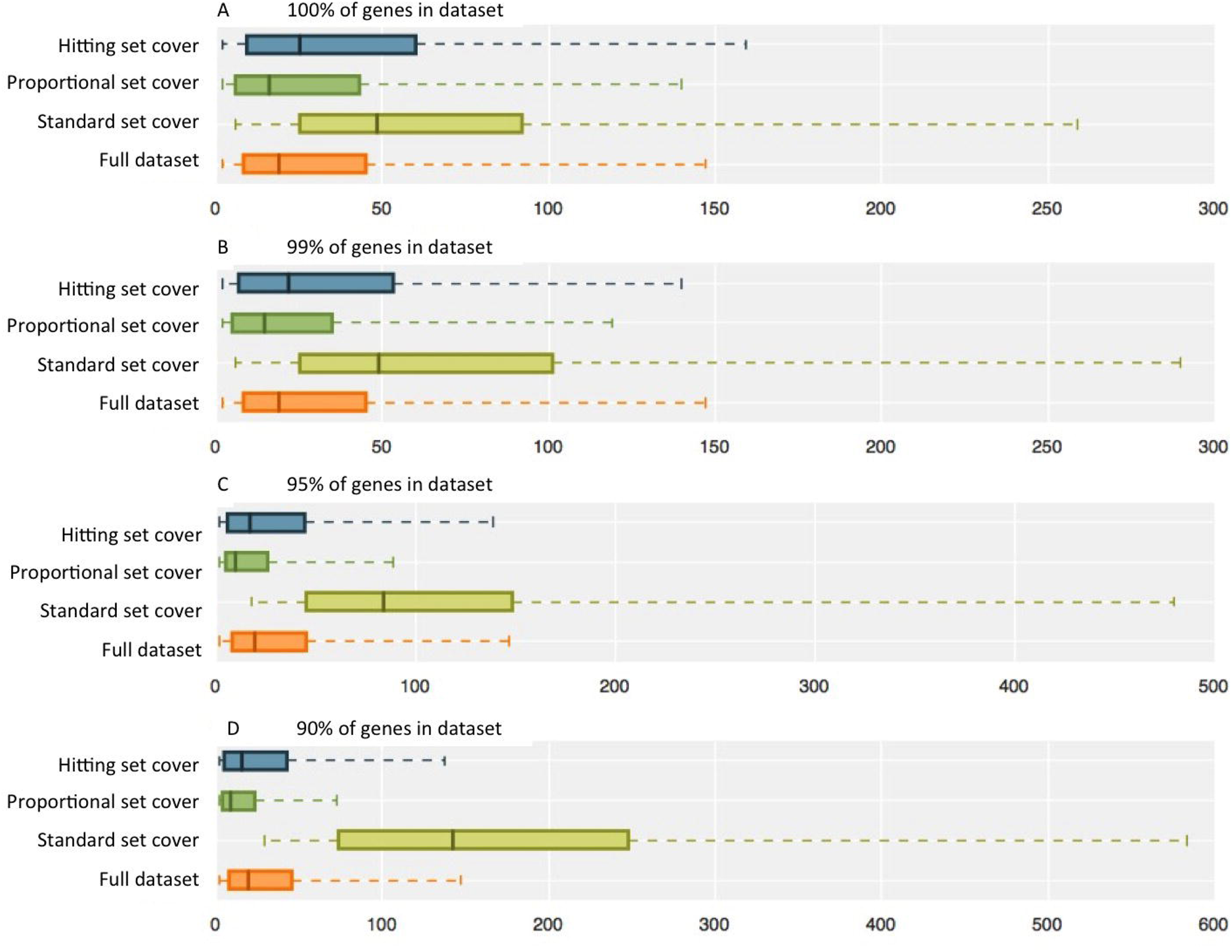
Pathway redundancy heat maps. (A) Pathway overlap for top ten enriched pathways. (B) Pathway overlap for top ten enriched pathways after application of set cover. The values represent asymmetric overlap, i.e. for each pathway shown on the left axis, values represent the proportion of genes that are also included in the pathway shown on the bottom axis.

Figure 5B shows overlap between the top ten pathways after application of the enrichment set cover algorithm. Because the differentially expressed genes covered by the ‘collagen biosynthesis and modifying enzymes’ pathway are a subset of those covered by the ‘extracellular matrix organization’ pathway, the ‘collagen biosynthesis and modifying enzymes’ pathway is removed from the cover set (Figure 5B). The second pathway in this list therefore becomes ‘GPCR signaling g alpha q’. The ‘collagen formation’ and ‘class b 2 secretin family receptors’ pathways are also removed because the differentially expressed genes they cover are additionally covered by the more enriched pathways ‘extracellular matrix organization’ and ‘signal transduction’ pathways (respectively). Additionally, ‘GPCR signaling pertussis toxin’ and ‘GPCR signaling cholera toxin’ are absent from the returned list, as all of their differentially expressed genes are found in ‘GPCR signaling g alpha q’ or ‘signal transduction’.

Some pathways in the enrichment set cover do still show high levels of overlap, for example ‘wnt signalling network’ is included despite 89% of its differentially expressed genes being covered by ‘signal transduction’. This is acceptable because ‘signal transduction’ is more highly enriched than ‘wnt signalling network’, yet the ‘wnt signalling network’ is worth including as it contains three differentially expressed genes that are not in ‘signal transduction’. If ‘wnt signalling network’ had been excluded then these genes would not have been described by the most significant pathway available to represent them. The unmodified top ten enriched pathways only cover 78.0% of the enriched genes. Using the set cover enrichment algorithm increases this figure to 85.2% without disrupting the pathway order given by the enrichment p-values.

## 6. Discussion and conclusion

We described algorithms suitable for reducing overlap in large pathway data sets allowing multiple databases to be amalgamated without excessive redundancy impeding the usefulness of the resource. Standard set cover is the best algorithm to reduce the number of pathways required to cover the data set, but significantly increases pathway size, which can be controlled by proportional set cover or hitting set cover. The proportional set cover is the best algorithm for controlling pathway size and the hitting set cover is the preferred choice for covering all of the genes in the dataset with minimal pathway redundancy. We showed that reducing the *GC* parameter allows further reductions in pathway redundancy; for example, if only 95% of the genes in the CPDB dataset were covered redundancy can be reduced by up to 88%. In addition reducing *GC* increases pathway size control when the proportional set cover and hitting set cover algorithms are used.

For pathway enrichment analysis we aimed to reduce redundancy while selecting the most significantly enriched pathways based on p-values. As an application we used the modified set cover algorithm to reduce the results of enrichment analysis from a large osteoarthritis data set. We found that 5 out of the 10 top ranking pathways could be omitted as they were subsets of more highly enriched pathways. Overlap between pathways returned from enrichment data is not always immediately obvious and requires further consideration. By reducing this redundancy, data interpretation is made more intuitive. Reducing redundancy also allows the user to explore substantially more of the data set using the same number of pathways.

The enrichment set cover algorithm presented within this study differs from existing methods implemented by ReCiPa and Pathcards, since enrichment analysis is performed prior to reduction of redundancy. This is because the different sets of pathway boundaries available in the full dataset may optimally fit the differentially expressed genes. For example, comparison of the ‘apoptosis’ taken from KEGG, Reactome and Wikipathways, reveals that many of the proteins are specific to a single database [22]. This is due to the vague definition of pathway boundaries, as well as differing experimental focus on cellular contexts, such as tissues or disease states. Following enrichment analysis the pathways that are most significantly enriched are selected to represent the differentially expressed genes and superfluous pathways are removed. This prevents the top results from being dominated by large numbers of highly similar pathways.

Set cover uses greedy heuristic methods, which provide good approximations of the optimal solution in a time effective manner. These methods are extremely efficient and can be run in a matter of minutes, however it should be noted that they do not guarantee an optimal solution. This is particularly true for the proportional set cover algorithm where the randomness of early selections influences the result. However, all possible outcomes result in reduced redundancy. The enrichment set cover algorithm is exempt from these considerations unless multiple pathways have identical p-values.

We have provided a method to dramatically reduce redundancy in pathways facilitating a more concise portrayal of cellular processes, while avoiding the issues introduced by pathway merging. Our algorithms are publicly available and have wide applicability to analysis of pathway datasets from any organism.

## 7. List of abbreviations

CPDB: Consensus PathwayDB
GC: Gene cover
SNP: Single nucleotide polymorphism

## 8. Declarations

### Ethics approval and consent to participate

NA

### Consent for publication

NA

### Availability of data and materials

https://github.com/RuthStoney/set-cover-and-set-packing-to-reduce-redundancy-in-pathway-data

### Author contributions

All authors contributed to the design of the study. R.S. performed the analysis and wrote the manuscript. All authors edited the manuscript and approved its final version.

## Acknowledgements

We would like to thank Jamie Soul and Sara Dunn from the University of Manchester, for providing the osteoarthritis data.

## 11. Additional material

Supplementary table 1: Enriched pathways from the osteoarthritis dataset (p-value<0.05). The set cover column indicated the 23 pathways that were included in the set cover. Found in additional file 1.docx.

